## Supplemental Information and Figures for "TANK binding kinase 1 promotes BACH1 degradation through both phosphorylation-dependent and -independent mechanisms without relying on heme and FBXO22": BACH1_TBK1_Liu_20240216_SupplementaryInformation-Figures.pdf

**Title**

Supplemental materials and methods

Supplemental references

Supplemental figure legends

### Supplemental materials and methods

#### Cell culture

Human embryonic kidney 293T (HEK-293T) were maintained in DMEM-low glucose supplemented with 10% heat-inactivated FBS (Sigma Aldrich) and 0.1 mg/mL penicillin/streptomycin (Gibco). The cells used were limited to less than 20 passages.

#### Western blotting

The supernatant containing the proteins was fractionated on slab gels (7.5% or 10% acrylamide separating gel; 4% stacking gel). The gels were run at 100V for 2 hours, and wet-transferred onto polyvinylidene difluoride (PVDF) at 300mA for 1.5 hours at 4 °C. Blots were washed with TBS and blocked for 1 hour in TBS-T/skimmed milk. The primary antibodies were diluted with 5% nonfat dried milk in TBS-T, the membranes were incubated at 4°C for one night. Washes (3 times, 10 min each in TBS-T) were done after primary and secondary antibody incubations. Secondary antibody incubation was performed for 1 hour at RT in 5% nonfat milk in TBS-T. ECL was performed with Clarity Western ECL substrate (Bio-Rad).

#### Antibodies

Anti-BACH1 mAb (clone 9D11) was purified from the culture supernatant of selected hybridoma. Anti-BACH1 antiserum (A1-6) was reported previously(1). Other antibodies were ACTB (GTX109639, GeneTeX), E-cadherin (ab1416, Abcam), ferritin light chain (sc-74513,Santa Cruz Biotechnology), ferritin heavy chain (sc-376594, Santa Cruz Biotechnology), HO-1 (ADI-SPA-896, Enzo Life Sciences), Anti-DDDDKtag(M185-3L, MBL Life Science) and Ubiquitin (MFK-003, Nippon Bio-Test Laboratories Inc).

**Mass spectrometry for protein identification**

MEL cells were plated at approximately 1 x $10^6$ cells per well in 12-well plates and infected with control (FLAG-IL2R) or mBACH1-FLAG-IL2R-expressing retrovirus. ReCLIP was performed using virus-infected MEL cells selected by the antibody of IL2R. Approximately $1 \times 10^8$ cells were fixed in PBS containing 0.5 mM DSP and DTME, then lysed in 1 ml of RIPA buffer. Samples were immunoprecipitated using anti-FLAG beads, followed by mass spectrometry. MASCOT was used for protein identification and carried out as described previously (2).

#### Supplemental figure legends

**Supplementary Fig. S1. FBXO22 promotes BACH1 degradation.**

**A,** MEL cells were infected with control (FLAG-IL2R) or mBACH1-Flag-IL2R-expressing retrovirus. ReCLIP was performed using virus-infected MEL cells selected by the antibody of IL2R. Samples were immunoprecipitated using anti-FLAG beads, followed by mass spectrometry. MASCOT was used for protein identification.

**B**, Namalwa cells were transfected with siRNAs for control (siC) or FBXO22 (siFBXO22) and treated with cycloheximide and hemin for indicated period. Western blots with indicated antibodies are shown.

**Supplementary Fig. S2. TBK1 promotes BACH1 degradation.**

**A and B,** HEK293T cells were transfected as written above and then were incubated with CQ (100 μM) or MG132 (10 μM) for 12 hours.

**C,** HEK293T cells were transfected with the indicated plasmids for 48 hours and were incubated with CQ (100 μM) or MG132 (10 μM) for the last 12 hours. l.ex., long exposure; s.ex., short exposure.

**Supplementary Fig. S3. Schematic representations of Plasmids.**

A

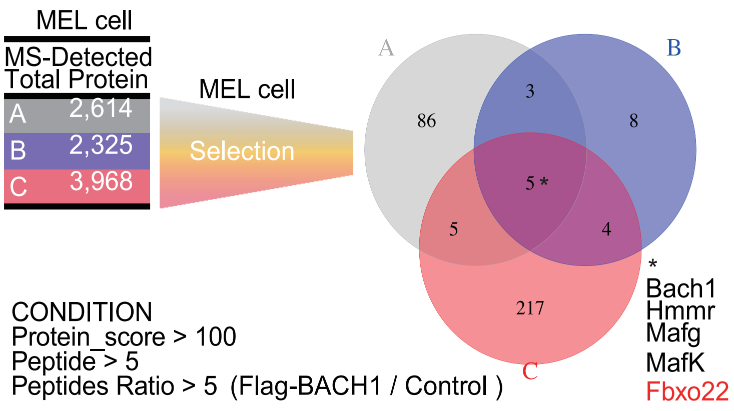
MEL cell
MS-Detected Total Protein
A 2,614
B 2,325
C 3,968
MEL cell
Selection
A
B
C
86
3
8
5 *
5
4
217
*
Bach1
Hmmr
Mafg
MafK
Fbxo22
CONDITION
Protein_score > 100
Peptide > 5
Peptides Ratio > 5 (Flag-BACH1 / Control )

B

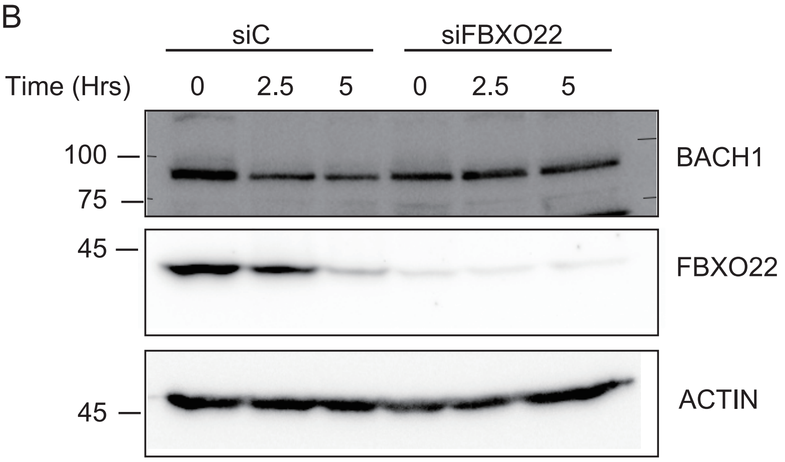
siC
siFBXO22
Time (Hrs)
0
2.5
5
0
2.5
5
100
75
BACH1
45
FBXO22
45
ACTIN

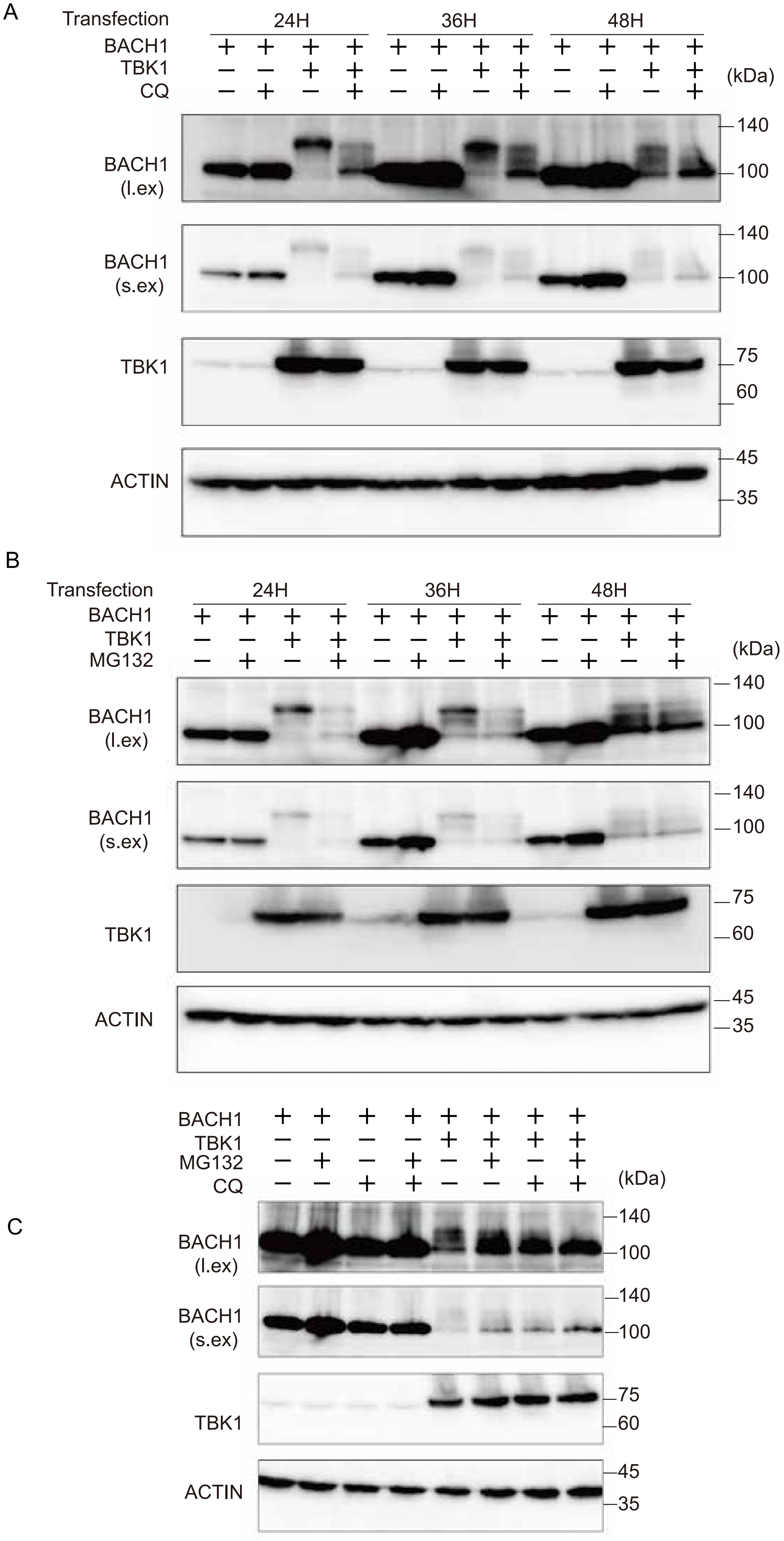
A
Transfection
24H
36H
48H
BACH1
TBK1
CQ
(kDa)
BACH1 (l.ex)
BACH1 (s.ex)
TBK1
ACTIN
140
100
75
60
45
35
B
Transfection
24H
36H
48H
BACH1
TBK1
MG132
(kDa)
BACH1 (l.ex)
BACH1 (s.ex)
TBK1
ACTIN
140
100
75
60
45
35
C
BACH1
TBK1
MG132
CQ
(kDa)
BACH1 (l.ex)
BACH1 (s.ex)
TBK1
ACTIN
140
100
75
60
45
35

Supplementary Fig. S3

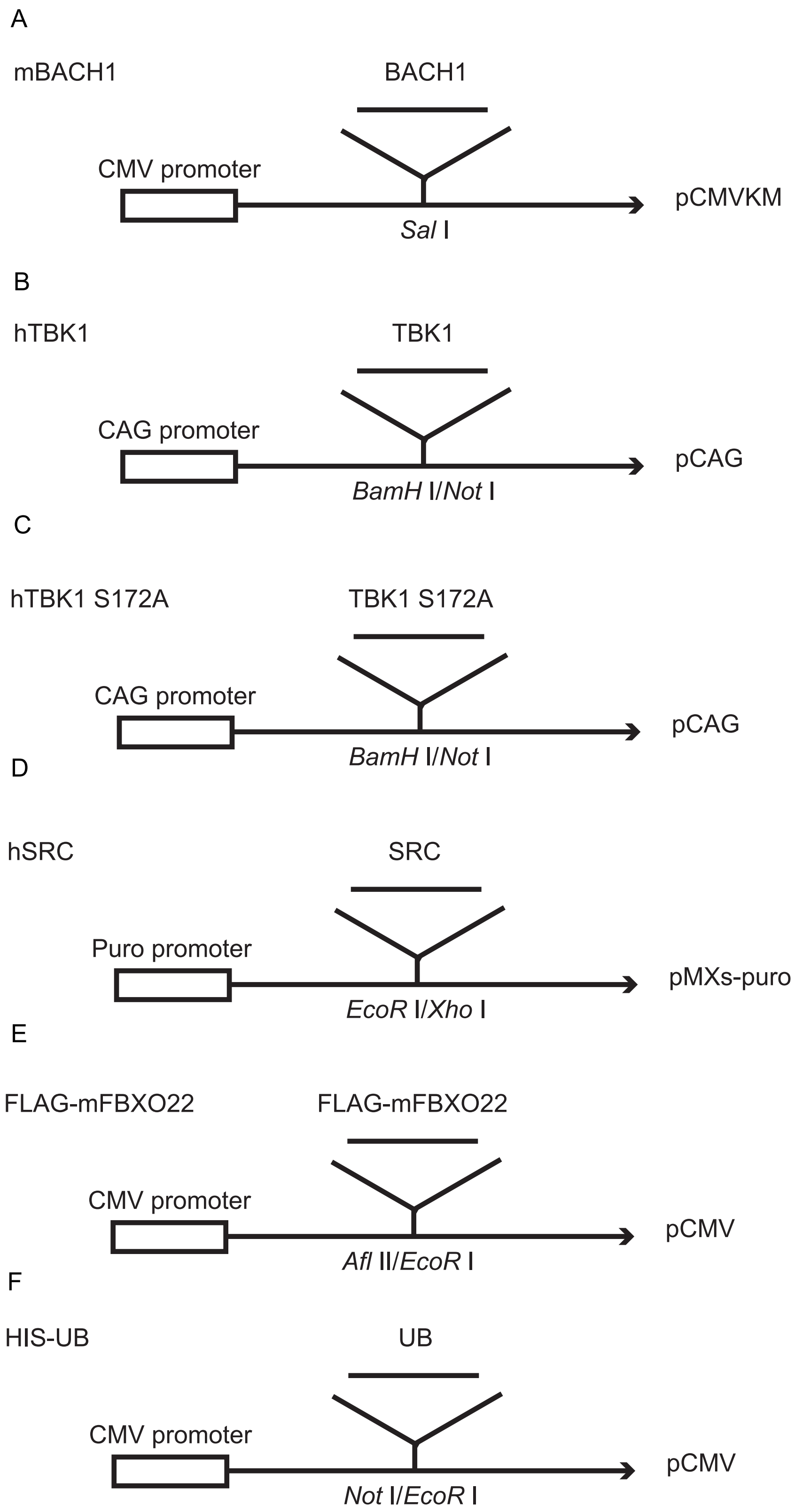
